# Structural basis of phosphate export by human XPR1

**DOI:** 10.1101/2024.12.30.630499

**Authors:** Yifei Wang, Yuechan Wang, Hui Yang, Ao Li, Dan Ma, Huaizong Shen

## Abstract

Phosphate is essential for all life forms because of its indispensable roles in energy metabolism, nucleic acid and phospholipid synthesis, cellular signaling, and the formation of bones and teeth. Its homeostasis is maintained through balanced import and export processes. In humans, XPR1 has been proposed as the sole phosphate exporter, although some controversy remains. Here, we report the closed and open structures of human XPR1, purified in the absence and presence of exogenous InsP6, respectively. The exporter forms a symmetric homodimer, with transmembrane helix (TM) 1 serving as the dimeric interface. The transmembrane domain of each protomer contains a transport module (TM5-10) and a supporting module (TM1-4). Compared to the closed XPR1, the open structure demonstrates a significant displacement of the extracellular portion of TM9 towards the periphery of the phosphate transport pathway, creating an open portal to the extracellular milieu. Additionally, a potential phosphate ion coordination site is strategically located in the middle of the transport pathway. On the intracellular side, the pathway entrance is obstructed by a cluster of residues, the C-plug, positioned on the carboxyl side of TM10. Consistently, the removal of the C-terminus significantly increases the transport activity of XPR1 in reconstituted liposomes. Our findings provide a comprehensive understanding of the export mechanism of human XPR1 and hold promise for the development of potential therapeutics for primary familial brain calcification and ovarian cancer.

---

Inorganic phosphate (Pi) is an essential macronutrient for life, playing crucial roles in energy metabolism, macromolecule synthesis, cellular signaling, and the formation of bones and teeth. Its homeostasis is orchestrated through a finely tuned equilibrium of import and export processes^1^. In humans, two Na+-dependent phosphate importer families, SLC34 and SLC20, have been established to be instrumental in phosphate import.

Conversely, our understanding of the phosphate export process remains limited. The only phosphate exporter proposed thus far in humans is XPR1, also denoted as SLC53A1. XPR1 was initially identified as a cell surface receptor for xenotropic and polytropic mouse leukemia viruses^2-4^. Later investigations suggest that XPR1 might function as a phosphate exporter, as evidenced by cell-based assays^5^. XPR1 mutations are linked to diseases including Primary Familial Brain Calcification (PFBC), hypophosphatemic rickets, and severe renal tubular dysfunction, possibly due to disturbance in cellular phosphate homeostasis^6,7^. Recent research has underscored its pivotal role in the survival of certain cancers such as Ovarian Clear Cell Carcinoma (OCCC)^8^.

XPR1 comprises three domains: the amino-terminal (N-terminal) soluble SPX domain (named after proteins Syg1, Pho81, and XPR1), the transmembrane domain, and the carboxyl-terminal (C-terminal) domain^9^. The SPX domain, which is implicated in several phosphate metabolism pathways, can bind to and be regulated by inositol polyphosphates (IPs) including inositol hexakisphosphate (InsP6), 5-diphosphoinositol 1,2,3,4,6-pentakisphosphate (5PP-InsP5 or InsP7), and 1,5-bisdiphosphoinositol 2,3,4,6-tetrakisphosphate (InsP8)^10^. The transmembrane domain of XPR1 is predicted to encompass ten transmembrane helices. The final few transmembrane helices comprise the EXS domain (named after proteins Erd1, XPR1, and Syg1) and are postulated to form the phosphate ion transport pathway^11^. Although the C-terminal domain of XPR1 is predicted to be unstructured, it is suggested to play critical regulatory roles in phosphate transport, potentially through interplay with SLC34 or SLC20 importers, albeit through unknown molecular mechanisms^9,12^.

To further explore the phosphate export mechanism of XPR1, we have determined its high-resolution structures in both closed and open states using the electron cryogenic microscopy (cryo-EM) method (Fig. 1a,b). Building upon our structural observations, the transport activity of strategically designed XPR1 constructs was examined using the phosphate efflux assay in HEK293T cells (see Materials and methods).

**Fig. 1.**
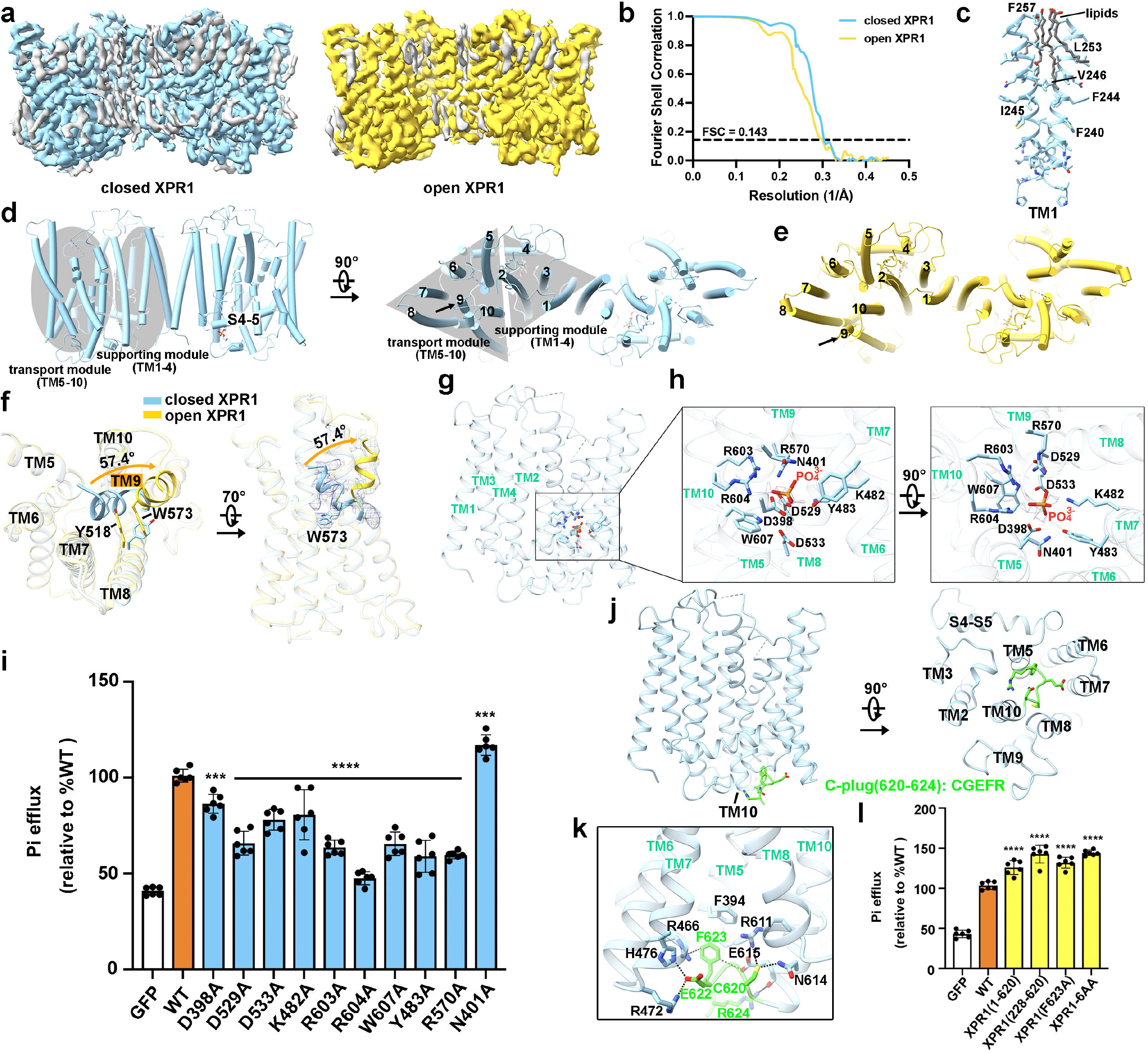
Structural basis of phosphate transport by human XPR1. **(a)** The cryo-EM density maps of XPR1 in closed (left) and open (right) states. **(b)** The Fourier shell correlation (FSC) curves for the cryo-EM reconstructions of XPR1 in closed (blue) and open states (yellow) indicate a resolution of 3.30 Å and 3.33 Å, respectively. **(c)** The dimeric interface encompasses transmembrane helix (TM) 1 and surrounding lipids from both protomers which engage in extensive hydrophobic interactions. **(d)** The closed structure of XPR1. The transmembrane domain of an XPR1 protomer comprises two relatively independent modules. TM1-4 constitute the supporting module, while TM5-10 form the transport module. **(e)** The open structure of XPR1. An evident difference between the closed **(d)** and open XPR1 **(e)** occurs in the extracellular portion of TM9, indicated by black arrows. **(f)** TM9 displacement between closed and open XPR1. From the closed to open XPR1, the extracellular portion of TM9 relocates from the center of the transport module to the periphery at a degree of 57.4°. Tyr518 on TM8 and Trp573 on TM9, which are distant in the closed structure, form π-π interactions that stabilize the open state. **(g)** A coordination site for the phosphate ion is strategically positioned at the center of the transport module, surrounded by residues from TM5-10. **(h)** The phosphate ion forms extensive electrostatic and H-bond interactions with these coordinating residues. (i) The roles of Pi-coordination residues were assessed through the Pi efflux assay. (j) Residues from 620 to 624 on the carboxyl side of TM10 constitute the C-terminal plug or C-plug. The C-plug binds to the intracellular exit of the phosphate transport pathway, enclosed by TM5-10. **(k)** The C-plug of XPR1 forms extensive interactions, including H-bonds, π-π, and cation-π interactions, with residues from the surrounding transmembrane helices. **(l)** XPR1 constructs with deleted C-terminus or mutations in the C-plug were assayed with the Pi efflux assay. For panels **(i)** and **(l)**, the activity of cells expressing GFP was used as control, and the activity of all mutants were normalized to the wild-type XPR1. The results are presented as mean ± s.e.m., with n = 6 from three independent experiments. Statistical analysis was performed using one-way ANOVA, with significance levels indicated as ^***^p < 0.001 and ^****^p < 0.0001.

The export activity of the full-length human XPR1 (UniProt ID: Q9UBH6), tagged with C-terminal GFP and FLAG tags, was first examined with the phosphate efflux assay (Fig. 1i,l, and Fig. S7). Compared to the cells expressing GFP as control, cells transfected with the tagged XPR1 display a significantly elevated phosphate efflux activity. The same construct was recombinantly expressed in HEK293F cells for structural determination. The target proteins were purified by anti-FLAG-tag G1 antibody and size-exclusion chromatography (SEC) sequentially. Probably due to the incomplete loss of the GFP tag during purification, two protein bands corresponding to XPR1 were visualized in the SDS-PAGE gel (Fig. S1a). Peak fractions of XPR1 from SEC were pooled together and utilized directly for structural determination of the closed XPR1. To elucidate the open structure, 1mM of InsP6, a commercially available substitute for PP-IPs, was supplemented throughout the purification process (Fig. S1b). Following a standard data acquisition and processing procedure, we determined the cryo-EM structures of closed and open human XPR1 at resolutions of 3.30 Å and 3.33 Å, respectively (Fig. 1a,b, Fig. S2, and Table S1). Only the transmembrane region of XPR1 was modeled which was determined at high resolution (Figs. S3, S4, and Table S1). In comparison, both the amino- and carboxy-termini are of relatively low resolution, indicating their flexibility. For an in-depth understanding of our structural determination and modeling details, please refer to the Materials and methods section.

The structure of XPR1 resembles the predicted one, which has been reported to represent a unique fold in the SLC superfamily^13^. It assembles as a symmetric homodimer (Fig. 1d,e). Each protomer contains ten transmembrane helices, with TM5 and TM6 harboring a kink in the middle. TM1 from these two protomers interact with each other via hydrophobic residues, together with several surrounding lipids forming the dimeric interface (Fig. 1a,c). These lipids may stabilize the dimer formation and contribute to the overall structural integrity of the transporter, which warrants further investigation. Careful inspection of the structure reveals that the transmembrane domain of XPR1 comprises two relatively independent structural modules: a transport module encompassing TM5-10 and a supporting module composed of TM1-4 (Fig. 1d). These two modules are connected by a long interfacial helix between TM4 and TM5, designated as S4-5. A probable phosphate transport pathway is enclosed by the transport module helices, arranged in a counterclockwise fashion from TM5 to TM10 when viewed from the extracellular side.

Notably, the transport module (residues 391-618) coincides with the previously proposed EXS domain (residues 439-643) in sequence^11^. In each protomer, a bifurcated lipid was discovered intercalated between the transport and supporting modules (Fig. S5). To investigate the potential role of this lipid, surrounding residues were mutated to tryptophan, a bulky amino acid intended to block the lipid’s presence. Surprisingly, these mutations did not significantly decrease the phosphate efflux activity of XPR1 (Fig. S7b), casting doubt on the lipid’s functional significance in the transport mechanism.

Inositol pyrophosphates (PP-IPs) are known to augment XPR1’s phosphate export activity by binding to its SPX domain^14^. We have determined the closed and open structures of XPR1 in the absence and presence of InsP6, a commercially available substitute for PP-IPs, respectively (Fig. 1a,g). A striking distinction between these two structures lies in TM9 (Fig. 1d-f). In the closed configuration, the extracellular portion of TM9 is centrally located within the transport module, effectively obstructing the transport pathway. Conversely, the extracellular part of TM9 in the open structure undergoes a pronounced shift towards the pathway’s periphery, resulting in an expanded opening to the extracellular milieu (Fig. 1f and Fig. S6a). The transport pathway’s radius dilates from 1 Å to approximately 4 Å at the extracellular exit when transitioning from the closed to open state (Fig. S6b). The conformational transition brings Tyr518 in TM8 and Trp573 in TM9, which are distant in the closed structure, into proximity, forming π-π interactions that stabilize the open state (Fig. 1f). Mutation of W573A substantially abrogates XPR1’s phosphate efflux activity, underscoring its critical role in the normal function of the transporter (Fig. S7a).

The phosphate permeation pathway within XPR1 is enclosed by helices of the transport module, comprising TM5-10 (Fig. 1d,g). Detailed examination of this pathway reveals a potential phosphate ion coordination site at its center, formed by Asp398 and Asn401 in TM5; Lys482 and Tyr483 in TM7; Asp529 and Asp533 in TM8; Arg570 in TM9; and Arg603, Arg604, and Trp607 in TM10 (Fig. 1h). Although the density is not well-defined, a potential phosphate ion is modeled at the center of this coordination site to facilitate analysis of its interactions with surrounding residues, which include extensive electrostatic and hydrogen-bond interactions. In the closed conformation, this site is located directly beneath Trp573, a key residue involved in conformational changes (Fig. 1f). Consistent with their critical roles, the phosphate efflux activity is generally reduced in XPR1 constructs where these coordination residues are mutated to alanine, although the extent of reduction varies (Fig. 1i). The N401A mutation is an exception, showing an increase in export activity, the reason for which awaits further investigation. To rule out the impact of surface expression on these findings, the surface expression profiles of these mutants were examined and confirmed (Fig. S8).

Succeeding TM10, a cluster of residues (620 to 624) obstructs the phosphate transport pathway at its cytoplasmic entrance (Fig. 1j and Fig. S3b). This cluster is referred to as the C-terminal plug, or C-plug. The C-plug residues engage in close interactions with neighboring residues from TM5-8 and TM10 (Fig. 1k). Notably, Phe623 interacts with Phe394, Arg466, and Glu615, while Glu622 engages with Arg472 and His476. Cys620 and Arg624 also play significant roles in this network. To investigate the C-plug’s function, constructs were designed to remove this obstruction, including a C-terminus deletion, an F623A mutation, and a mutation replacing residues 619NCGEFR624 with six alanines (designated as XPR1-6AA). Consistent with its inhibitory role, deleting the C-plug or mutating these critical residues significantly increased the transporter’s efflux activity (Fig. 1l).

Despite decades of studies, the export mechanism of XPR1 remains elusive. In this study, we resolved the structures of human XPR1 in both closed and open states, enabling a comparative analysis of the associated conformational changes. Notably, we identified a potential phosphate ion coordination site situated midway along the transport pathway.

Furthermore, we discovered the presence of a C-plug, which obstructs the intracellular entrance of the transport pathway. The structural observations were corroborated by the phosphate efflux assays.

In the transition from the closed to the open state of XPR1, a key conformational change involves the extracellular segment of TM9 shifting from the center to the periphery of the transport pathway. This rearrangement is prompted by the inclusion of InsP6 molecules during protein purification. The SPX domain of XPR1, which binds and is regulated by IP molecules in vitro, likely plays a crucial role in this process. Based on these insights, we propose that the binding of IPs/PP-IPs to the SPX domain induces conformational changes in XPR1, which involves displacing the C-plug from the intracellular entrance of the pathway and allowing phosphate ions to enter. Subsequent fliping of the Trp573 side chain leads to the structural repositioning of the extracellular portion of TM9, resulting in the opening of the exporter. Nevertheless, these conformational changes should be interpreted with caution, as the open structure obtained here may not fully represent the truly activated state of XPR1 under the regulation of genuine PP-IPs.

In summary, our structural findings offer valuable insights into the phosphate export mechanisms of human XPR1, including the identification of the potential phosphate ion coordination site, the C-terminal plug, and the conformational changes coupled to the phosphate transport process. These insights pave the way for the development of innovative therapeutic interventions for PFBC and ovarian cancer in the future.

## Data availability

Atomic coordinates and EM maps of closed human XPR1 (PDB: 9j97; EMDB: EMD-61254) and open human XPR1 (PDB: 9j98; EMDB: EMD-61255) have been deposited in the Protein Data Bank (http://www.rcsb.org) and the Electron Microscopy Data Bank (https://www.ebi.ac.uk/pdbe/emdb/).

## Acknowledgements

We thank the Cryo-EM Facility and Supercomputer Center of Westlake University for providing data collection and computation support, respectively. This work was supported by the National Science Foundation of China (32122042 and 32071208 to H.S.), the Zhejiang Provincial Natural Science Foundation (DQ24C050001 to H.S.), and the Westlake Education Foundation (to H.S).

## Author information

These authors contributed equally: Yifei Wang, Yuechan Wang, and Hui Yang.

## Authors and affiliations

Zhejiang Key Laboratory of Structural Biology, School of Life Sciences, Westlake University, Hangzhou, Zhejiang, China

Yifei Wang, Yuechan Wang, Hui Yang, Ao Li, Dan Ma & Huaizong Shen

Westlake Laboratory of Life Sciences and Biomedicine, Hangzhou, Zhejiang, China

Yifei Wang, Yuechan Wang, Hui Yang, Ao Li, Dan Ma & Huaizong Shen

Institute of Biology, Westlake Institute for Advanced Study, Hangzhou, Zhejiang, China

Yifei Wang, Yuechan Wang, Hui Yang, Ao Li, Dan Ma & Huaizong Shen

## Contributions

The project was conceived by H.S. Molecular cloning, phosphate efflux assay, protein expression profile examination, protein purification, cryo-sample preparation, and electron micrography data collection were performed by Yifei Wang, Hui Yang, and Ao Li.

Yuechan Wang conducted the cryo-EM map reconstruction and model building. All authors contributed to the data analysis. The manuscript was written by H.S.

## Corresponding authors

Correspondence to Huaizong Shen.

## Ethics declarations

**Competing interests**

The authors declare no competing interests.

