## Supplementary Information for "Structural basis of phosphate export by human XPR1"

### Materials and methods

#### Phosphate efflux assay of human XPR1 and mutants

The phosphate efflux assay was adapted from existing protocols with some modifications[1,2]. We seeded HEK293T cells (Gibco) at a density of $2 \times 10^5$ cells per well in 12-well plates, using DMEM medium (Gibco) supplemented with 10% FBS (WISENT). The cells were incubated at 37°C with 5% $CO_2$. After 24 hours, we transfected each well with 1 μg of plasmid DNA and 2 μl of Lipofectamine 3000 (Invitrogen). The assay was initiated 24 hours after transfection. Cells were washed three times with phosphate-free DMEM and incubated in the same medium at 37°C for 1 hour. We collected the supernatants and measured phosphate levels using the Malachite Green Phosphate Assay Kit (MAK307, Sigma). Cells were lysed with RIPA buffer (Beyotime) (50 mM Tris (pH 7.4), 150 mM NaCl, 1% Triton X-100, 1% sodium deoxycholate, 0.1% SDS, sodium orthovanadate, sodium fluoride, EDTA, and leupeptin), incubated at room temperature for 10 minutes, and then the lysates were transferred to 96-well plates for protein quantification using a BCA protein assay kit (Beyotime). Phosphate efflux activity was normalized to the total protein content. Each experiment was repeated at least three times on different days, with triplicate wells analyzed for each condition. The data were analyzed using Prism10 (Graphpad). To confirm plasma membrane localization, we used a Nikon C2Si inverted confocal microscope to examine HEK293T cells expressing GFP alone, GFP-tagged wild-type XPR1, and all mutants (Fig. S8).

#### Cloning, expression, and purification of human XPR1

The gene encoding human XPR1 (UniProt: Q9UBH6) was synthesized by GENEWIZ and confirmed by sequencing. Full-length coding sequence was cloned into a pEG BacMam expression vector with a C-terminal GFP and Flag tag in XPR1[3]. All the plasmids for transient expression were confirmed through DNA sequencing. Expi293F cells (Gibco, Thermo Fisher Scientific) were cultured in SMM 293T-II medium (Sino Biological) at 37 °C under 5% CO2 in a ZCZY-CS9 shaker at 150 rpm (Zhichu Instrument). When the cell density reached $2 \times 10^6$ cells per mL, 2mg plasmids of XPR1 were mixed with 6 mg of linear polyethylenimine (Yeasen) before being added to one liter of cells. Transfected cells were harvested after ~48 hours of culturing and frozen in liquid nitrogen before being stored in a −80 °C refrigerator for future use.

For the purification of XPR1, frozen cell pellets were resuspended in the lysis buffer (50 mM Tris-HCl pH 7.4, 150 mM NaCl, 1% DDM:CHS (10:1, w/w) (DDM and CHS were purchased from Anatrace Products), 1 μM leupeptin, 1 μM pepstatin A, 1 μg/mL aprotinin, 1 mM PMSF) for 2 hours at 4 °C with gentle agitation. Solubilized cell lysis was clarified by centrifugation at 13,000 rpm for one hour at 4 °C. The resulting supernatant was applied to Anti-DYKDDDDK G1 Affinity Resin (Nanjing GenScript Biotechnology) before being washed with 100 mL wash buffer (50 mM Tris-HCl pH 7.4, 150 mM NaCl, 1 mM PMSF, 0.03% GDN (Anatrace Products)). The target protein was eluted from the resin using 20 mL elution buffer (50 mM Tris-HCl pH 7.4,150 mM NaCl, 1 mM PMSF, 0.03% GDN, 200 μg/mL FLAG peptide (Nanjing GenScript Biotechnology)). Then the eluates were concentrated to 1 mL using an Amicon Ultra-4 centrifugal filter (MWCO 50 kDa) (Millipore, Sigma Aldrich Trading Co.) at 4 °C before being injected into a Supedex 200 10/300 Column (GE Healthcare). The SEC was performed in a buffer containing 50 mM

Tris-HCl pH 7.4, 150 mM NaCl, 1 mM PMSF, and 0.03% GDN. The peak fractions were collected and concentrated for cryo-EM sample preparation. The entire purification procedure for XPR1 with InsP6 was identical except an addition of 1mM InsP6 (Sigma, catalog #P8810) and 10mM $NaH_2PO_4$/$Na_2HPO_4$ throughout the process.

**Cryo-EM sample preparation and data acquisition**

To prepare cryo-EM sample, the concentrated protein with or without 1mM InsP6 added was placed on glow-discharged holey carbon grids (Quantifoil Au R1.2/1.3) and flash-frozen in liquid ethane cooled by liquid nitrogen using Vitrobot (Mark IV, Thermo Fisher Scientific). The grids were loaded onto a 300 kV Titan Krios equipped with a K3 Summit detector (Gatan) and a GIF Quantum energy filter. Images were automatically collected using AutoEMation[4] in super-resolution mode at a nominal magnification of 81,000×, with a slit width of 20 eV on the energy filter. A defocus series ranging from -1.5 μm to -2.0 μm was used. Each stack was exposed for 2.56 s with an exposure time of 0.08 s per frame, resulting in a total of 32 frames per stack. The total dose was approximately 50 $e^-$/Å2 for each stack. The stacks were motion-corrected with MotionCor2[5] and binned 2 fold, resulting in a pixel size of 1.0773 Å/pixel. Meanwhile, dose weighting was performed[6]. The defocus values were estimated with Gctf [7].

**Image processing**

A diagram for the data processing is presented in Figure S2. A total of 5,865 and 3,816 micrographs were collected for the closed and open XPR1, respectively. All the micrographs were imported to CryoSPARC v4. A total of 6,066,873 and 4,006,819 particles were autopicked by CryoSPARC for the closed and open XPR1, respectively. After several rounds of two-dimensional classification, 313,305 and 46,087 selected particles from 2D classification for the closed and open XPR1, respectively, were subjected to ab initio reconstruction, homogeneous refinement, and several rounds of heterogeneous refinement sequentially. Particles from the best class of the final round of heterogeneous refinement were subjected to non-uniform refinement, resulting in 3D maps with overall resolutions of 3.30 and 3.33Å for the closed and open XPR1, respectively.

**Model building and structure refinement**

The predicted model for human XPR1 (AlphaFold Protein Structure Database ID: Q9UBH6)[8,9] was docked into the final maps of the closed and open XPR1 in ChimeraX[10]. Every residue was manually checked and adjusted in COOT[11]. The chemical properties of amino acid residues were considered during model building. The N terminus (residues 1–227), two residues of S4-5 linker (residues 435, 436), and the C terminus (residues 627-696) are invisible for both the closed and open XPR1 structures; hence, these regions were not built in the models. The extracellular half of TM9 and the S9-10 linker in the closed XPR1 are of low resolution; hence the residues 581-584 were not built. Two $PO_4^{3-}$ ions were tentatively modeled in the transport pathways for the closed XPR1 structure with the chemical environment considered. Several lipid molecules were tentatively modeled based on the map densities. Structure refinement was performed using the phenix.real_space_refine application in PHENIX in real space with secondary structure and geometry restraints[12]. The overfitting of the overall model was monitored by refining the model in one of these two independent maps from the gold-standard refinement approach and testing the refined model against the other map[13]. Statistics of the map reconstruction and model refinement can be found in Table S1.

**Preparation of liposome and proteoliposome**

DOPC lipids (Avanti Polar Lipids) were dissolved in chloroform/methanol mixture (3:1, v/v) at 50 mg $ml^{-1}$ and dried using nitrogen gas. To make preformed liposomes, lipids were resuspended at 20 mg/ml in a buffer containing 20 mM HEPES (pH 7.0) before being frozen and thawed ten times using liquid nitrogen. For reconstitution, liposomes were extruded through 0.1 μm membrane filters (Whatman) and incubated with 1% n-octyl-β-d-glucoside (β-OG; Anatrace) for 30 minutes at 4°C. Detergent-solubilized proteins were added into liposome/β-OG mixture at a protein: lipid ratio of 1: 100 (w/w) before being incubated for 2 hours at 4°C. Prewashed Bio-Beads SM2 (BioRad) was added to remove detergent (1g Bio-Beads for 117 mg OG). The amount of Bio-beads calculated above were added into the mixtrue for four times and incubated at 4°C: the first time for 1 hour, the second time for 1 hour, the third time overnight, and the fourth time for 1 hour. The proteoliposomes were harvested by ultracentrifugation at 28,000 rpm for 1 hour the next day. Then the proteoliposomes were resuspended with a second buffer (20 mM HEPES, pH 7.0) to a final lipid concentration of 5 mg/ml for the phosphate permeability assay.

**Stopped-flow assay for phosphate permeability**

To measure the phosphate permeability, the reconstituted liposomes or proteoliposomes were rapidly mixed with a hypertonic buffer containing 200 mM $Na_2HPO_4$/ $NaH_2PO_4$ at a volume ratio of 1:1. Alteration of vesicle size was detected by recording the light scattering signal at 440 nm. A stopped-flow spectrometer (Applied Photophysics SX 20) was used for the test at room temperature. The recorded data were analyzed using Prism10 (Graphpad).

### Supplementary figures and tables

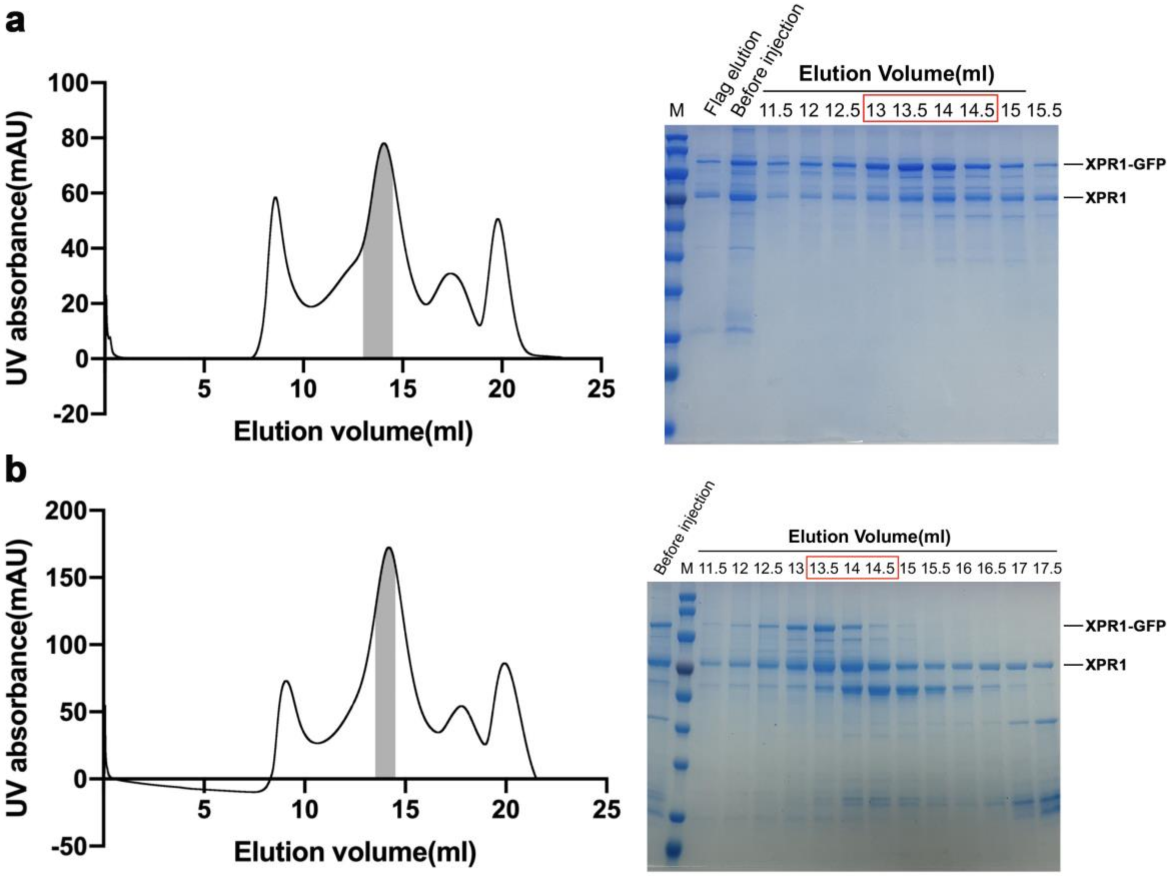


**Fig. S1 | Size-exclusion chromatography (SEC) profile and SDS-PAGE gel of human XPR1.** (**a**)(**b**) Size exclusion chromatography (SEC) profile and SDS-PAGE gel for preparations of closed (**a**) and open (**b**) human XPR1 are displayed. The peak fractions for the final sample preparations are depicted in gray in the SEC results and boxed in the corresponding SDS-PAGE gels stained with coomassie blue.

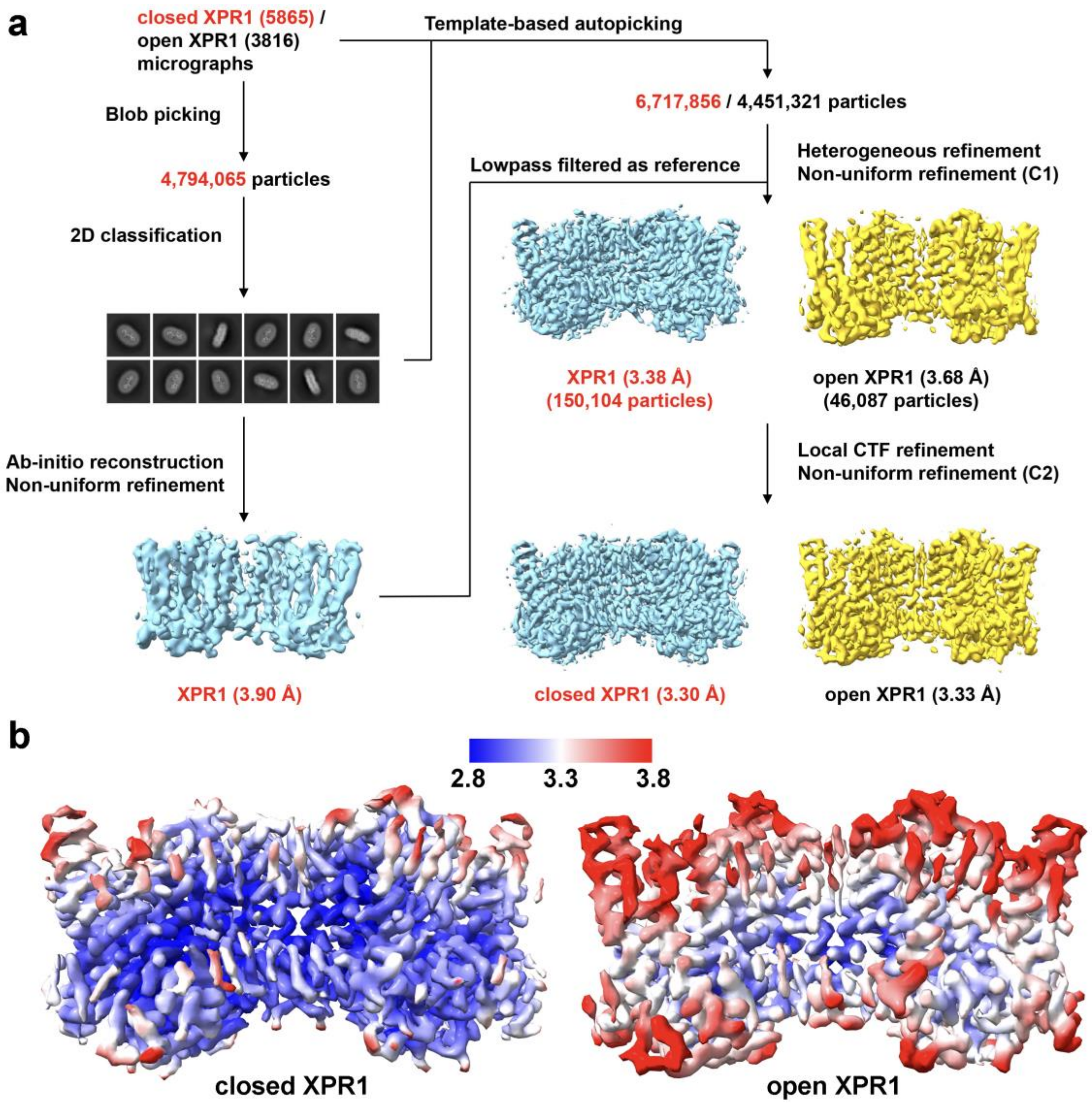


**Fig. S2 | Cryo-EM data processing flowchart for closed and open XPR1.** (**a**) Cryo-EM data processing flowcharts for both closed and open XPR1 are illustrated in detail. Representative 2D and 3D results are illustrated with available resolutions for 3D reconstructions indicated. Data processing was performed in CryoSPARC[14]. (**b**) Local resolution maps of closed (left panel) and open (right panel) XPR1 were calculated using CryoSPARC and presented in ChimeraX.

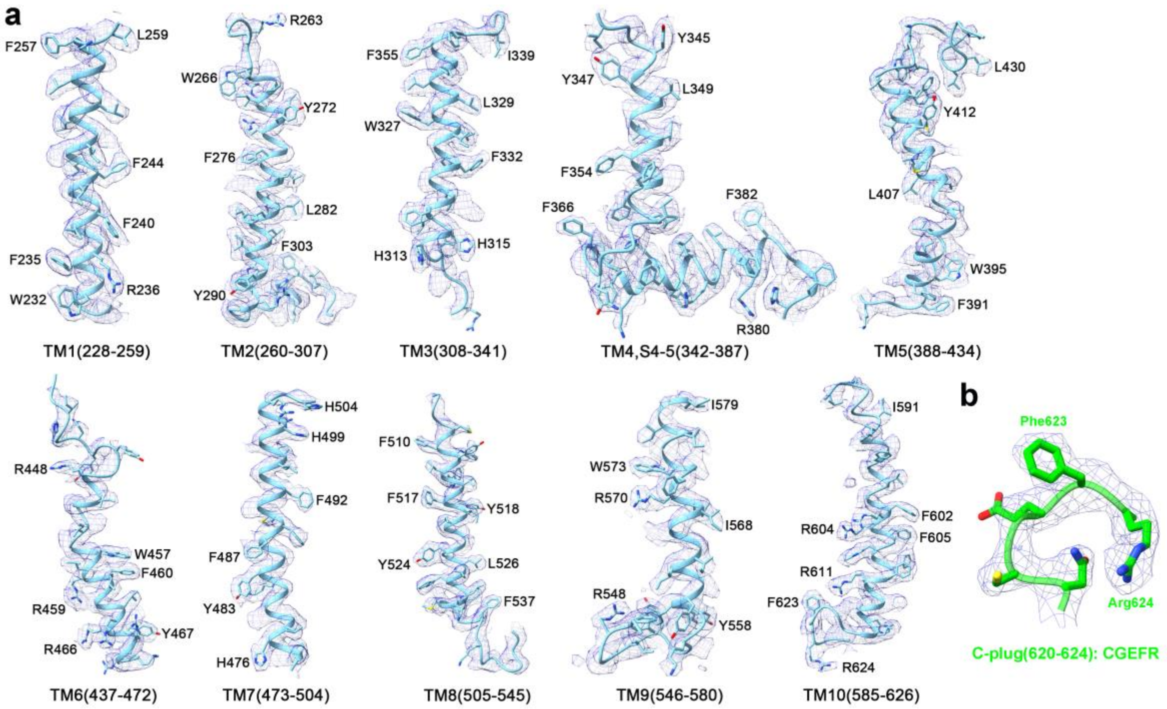


**Fig. S3 | EM densities for the transmembrane segments of closed XPR1.** (**a**)(**b**) The electron microscopy (EM) densities for the transmembrane segments (**a**) and the C-plug (**b**) of the closed XPR1 structure are visualized using UCSF ChimeraX[10]. Residues with large side chains are labeled.

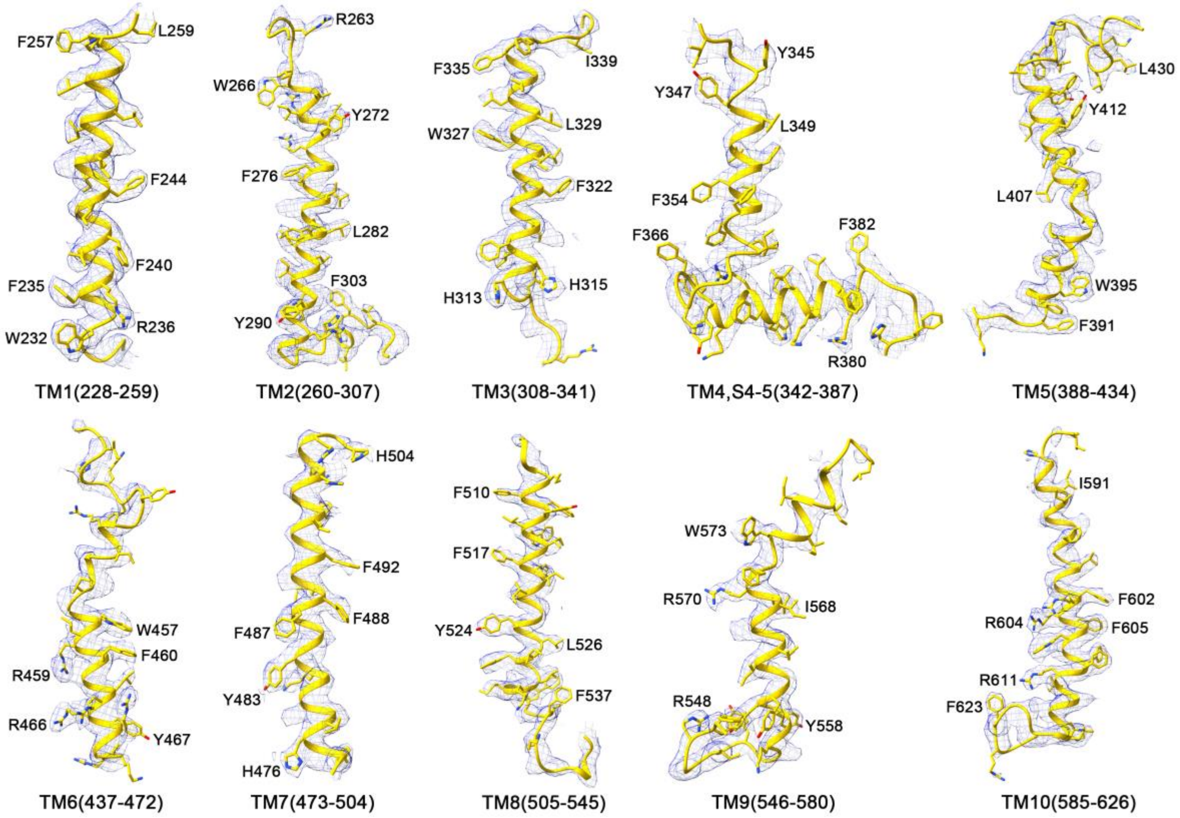


**Fig. S4 | EM densities for the transmembrane segments of closed XPR1.** The electron microscopy (EM) densities for the transmembrane segments of closed XPR1 structure are visualized using UCSF ChimeraX[10]. Residues with large side chains are labeled.

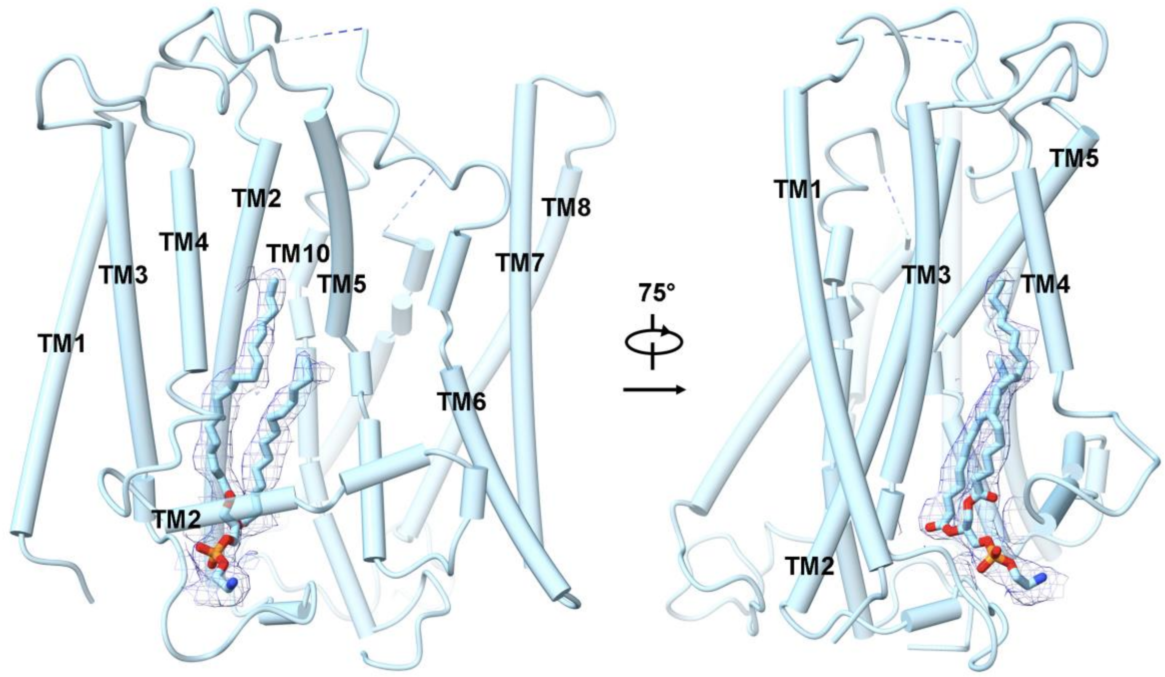


**Fig. S5 | An intercalated lipid.** A bifurcated lipid is intercalated between the transport and supporting modules at the membrane leaflet proximal to the cytoplasmic side. The tails of the lipid form hydrophobic contacts with surrounding transmembrane helices including TM2-5.

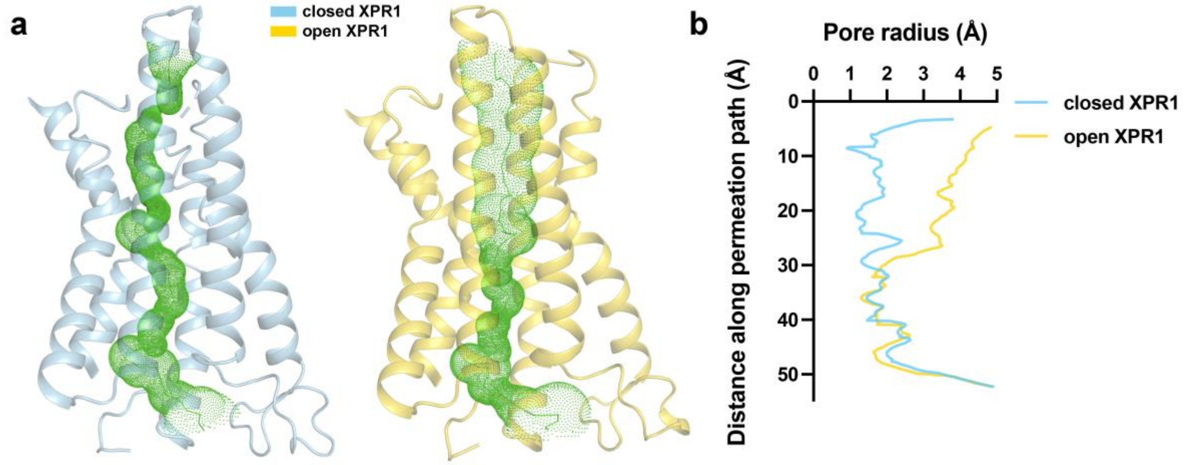


**Fig. S6 | Phosphate transport pathway comparison between closed and open XPR1.** (**a**) Phosphate transport pathways in closed and open XPR1. The pore near the extracellular exit of the pathway in open XPR1 is much wider than in the closed structure. The transport pathways were calculated by HOLE[15] and represented by green dots. The figure was prepared in PyMOL[16]. (**b**) Comparison of the pore radii. Considerable discrepancies of the pore radii near the extracellular side are observed between the transport pathways of XPR1 in closed and open states.

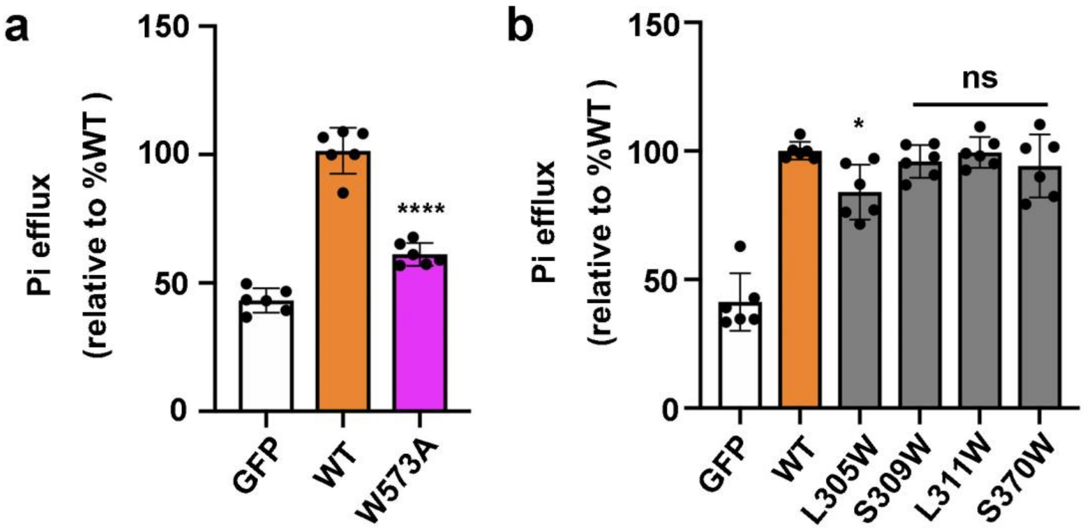


**Fig. S7 | Phosphate efflux assays on XPR1(W573A) and residues surrounding the intercalated phospholipid.** (**a**)(**b**) The function of Trp573 (**a**) and the residues surrounding the phospholipid (**b**) were assayed using the Pi efflux assay. The activity of cells expressing GFP was used as control, and that of all constructs were normalized to the wild-type XPR1. The results are presented as mean ± s.e.m., with n = 6 from three independent experiments. Statistical analysis was performed using one-way ANOVA, with significance levels denoted as $*p < 0.05$, $****p < 0.0001$, and "ns" indicating not significant.

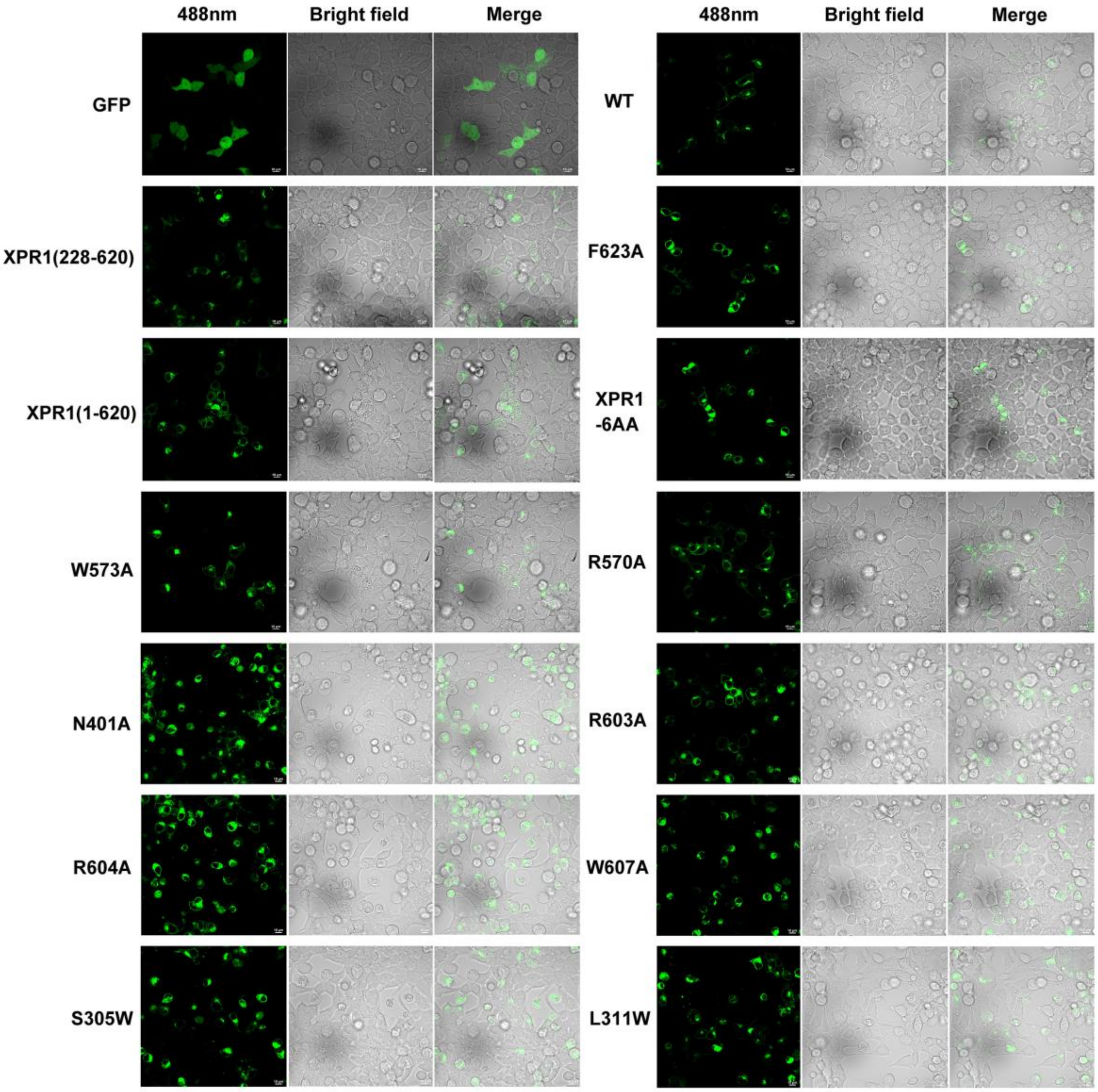


**Fig. S8 | Surface expression profiles of XPR1 and mutants.** The expression and distribution of XPR1 and its mutants were analyzed using fluorescence confocal microscopy, with GFP-expressing cells serving as the control. Both fluorescence and bright-field images were captured and merged for display.

**Table S1. Statistics for data collection and structural refinement.**

| **Data collection** | closed XPR1 | open XPR1 |
|---|---|---|
| EM equipment | Titan Krios (Thermo Fisher Scientific Inc.) | |
| Voltage (kV) | 300 | |
| Detector | Gatan K3 Summit | |
| Energy filter | Gatan GIF Quantum, 20 eV slit | |
| Pixel size (Å) | 1.087 | |
| Electron dose ($e^-$/Å$^2$) | 50 | |
| Defocus range (μm) | -1.5 ~ -2.0 | |
| Number of collected movie stacks | 5865 | 3816 |
| **Reconstruction** | | |
| Software | CryoSPARC v4 | |
| Number of used particles | 150,104 | 46,087 |
| Symmetry | C2 | C2 |
| Overall resolution (Å) | 3.30 | 3.33 |
| Map sharpening B-factor (Å$^2$) | -163.4 | -145.9 |
| **Refinement** | | |
| Software | Phenix | |
| Cell dimensions | | |
| a=b=c (Å) | 278.272 | |
| α=β=γ (˚) | 90 | |
| Model composition | | |
| Protein residues | 786 | 794 |
| Side chains assigned | 786 | 794 |
| $PO_4^{3-}$ | 2 | 0 |
| CLR | 2 | |
| 8PE | 2 | |
| R.m.s deviations | | |
| Bonds length (Å) | 0.006 | 0.005 |
| Bonds angle (˚) | 0.666 | 0.540 |
| Ramachandran plot statistics (%) | | |
| Preferred | 91.17% | 89.05% |
| Allowed | 8.57% | 10.70% |
| Outlier | 0.26% | 0.26% |
